# Optimization-Based Decoding of Imaging Spatial Transcriptomics Data

**DOI:** 10.1101/2022.11.22.517523

**Authors:** John P. Bryan, Loïc Binan, Cai McCann, Yonina C. Eldar, Samouil L. Farhi, Brian Cleary

## Abstract

**Motivation:** Imaging Spatial Transcriptomics (iST) techniques characterize gene expression in cells in their native context by imaging barcoded probes for mRNA with single molecule resolution. However, the need to acquire many rounds of high-magnification imaging data limits the throughput and impact of existing methods.

**Results:** We describe the Joint Sparse method for Imaging Transcriptomics (JSIT), an algorithm for decoding lower magnification IT data than that used in standard experimental workflows. JSIT incorporates codebook knowledge and sparsity assumptions into an optimization problem which is less reliant on well separated optical signals than current pipelines. Using experimental data obtained by performing Multiplexed Error-Robust Fluorescence in situ Hybridization (MERFISH) on tissue from mouse motor cortex, we demonstrate that JSIT enables improved throughput and recovery performance over standard decoding methods.

**Availability and Implementation:** *Software implementation of JSIT, together with example files, are available at* https://github.com/jpbryan13/JSIT.

**Supplementary Information:** Supplementary data are available at *Bioinformatics* online.

## 1 Introduction

Imaging Spatial Transcriptomics (iST) methods, such as MERFISH (K. H. Chen et al. 2015), CosMx (He et al. n.d.), and STARMAP (Wang et al. 2018) simultaneously measure expression of targeted sets of hundreds of genes at a time with single molecule spatial resolution, revealing spatial patterns of cell type arrangements and tissue organization. These newly developed methods have already allowed researchers to construct high-resolution spatial tissue atlases (Zhang et al. 2021), study subcellular compartmentalization of gene expression (Xia et al. 2019), and observe spatial differences in gene expression between phenotypical conditions (Moffitt, Bambah-Mukku, et al. 2018), with potential for further discovery as data sets grow and analysis frameworks mature.

All iST methods achieve gene multiplexing using combinatorial barcoding, in which each gene is assigned a distinct binary barcode from a pre-defined codebook (Tian, F. Chen, and Macosko 2022). Fluorescent probes complementary to the genes are then iteratively applied, imaged at high resolution, and removed from the sample, such that an individual mRNA molecule (or transcript) only appears as a bright spot in the images corresponding to the ones in its barcode. Software pipelines then perform computational decoding of the acquired images, identifying the location of each fluorescent spot and attempting to assign it to a gene identity. These decoded transcripts are then assigned to a cell, and all transcripts within a cell are tallied up to produce a count table relating cell location to gene expression.

Notably, while single molecule resolution imaging is necessary to identify transcripts, once the count table is produced, most downstream methods do not need individual molecule locations to perform common tasks such as cell type identification and differential gene expression analysis (Stogsdill et al. 2022; Dries et al. 2021; Hu et al. 2021). These analyses stand to be better empowered by profiling larger sample areas to increase the likelihood of capturing rare cell types and interactions and of profiling distinct regions of the tissue. However, the requirement of single molecule resolution imaging has thus far set the maximum imaging throughput to roughly 1 cm^2^ per day, limiting the biological discovery impact of iST. If, instead, imaging could be performed at lower magnification, larger amounts of tissue could be studied and iST could better enable the study of developmental time courses, comparisons among large numbers of patient samples, and the creation of large-scale tissue atlases.

In this work we re-frame the decoding problem as an optimization problem and leverage algorithmic techniques like those used in super-resolution microscopy (Solomon et al. 2018), to enable decoding of lower-magnification iST data. iST data is known to be structured: fluorescence signals are sparse in spatial coordinates. Currently used decoding pipelines, such as MERlin (Emanuel, Eichorn, and Sepulveda 2020), deconvolve optical signals and assign them to most likely barcodes in distinct steps, without taking full advantage of the known sparsity of the data. We instead present the Joint Sparse method for Imaging Transcriptomics (JSIT), which combines optical deconvolution and decoding of iST data, explicitly incorporating signal sparsity knowledge into the decoding step. In the process, this joint approach relaxes the requirement for high magnification imaging, and increases overall throughput.

In the remainder of the paper, we first provide a detailed description of the JSIT algorithm and problem formulation; its solution with the iterative proximal gradient algorithm FISTA (Beck and Teboulle 2009); and the metrics chosen to compare JSIT to MERlin. We then describe JSIT’s performance on real MERFISH data from the mouse primary motor cortex (MOp) imaged at both 40x and 60x. In particular, we investigate JSIT’s ability to perform cell typing on low-resolution imaging data. We show that using JSIT on 40x data recapitulates the cell typing of using MERlin on 60x data, while MERlin is not able to achieve this at 40x. Finally, we discuss the practical throughput advantages of using JSIT to decode MERFISH data.

## 2 Methods

### 2.1 Problem formulation: modeling iST data

We begin by modeling the iST data generation process, as depicted schematically in Fig. 1a. Our aim is to recover the spatial distribution and gene identity of mRNA from raw data consisting of *F* images with size *n* × *n* pixels, which encode gene identities with defined barcodes of *F* bits. We first vectorize and subsequently concatenate these images to form a matrix 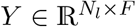, where *N*_*l*_ = *n*^2^ is the number of pixels in the field of view (FOV). Element (*i, f*) of *Y* represents the pixel value at location *i* in the *f* th image. We model the generation of *Y* as:

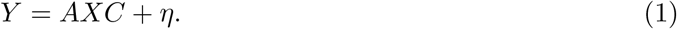

Here, *A* represents the operation of the PSF of the microscope, *X* represents the spatial distribution and gene identity of mRNA, which we aim to recover, *C* is the codebook, and *η* denotes additive experimental noise. This noise is not defined in specific terms, but is known to include low-frequency autofluorescence background, which varies strongly with location within the FOV, and high-frequency, uncorrelated shot noise.

**Figure 1:**
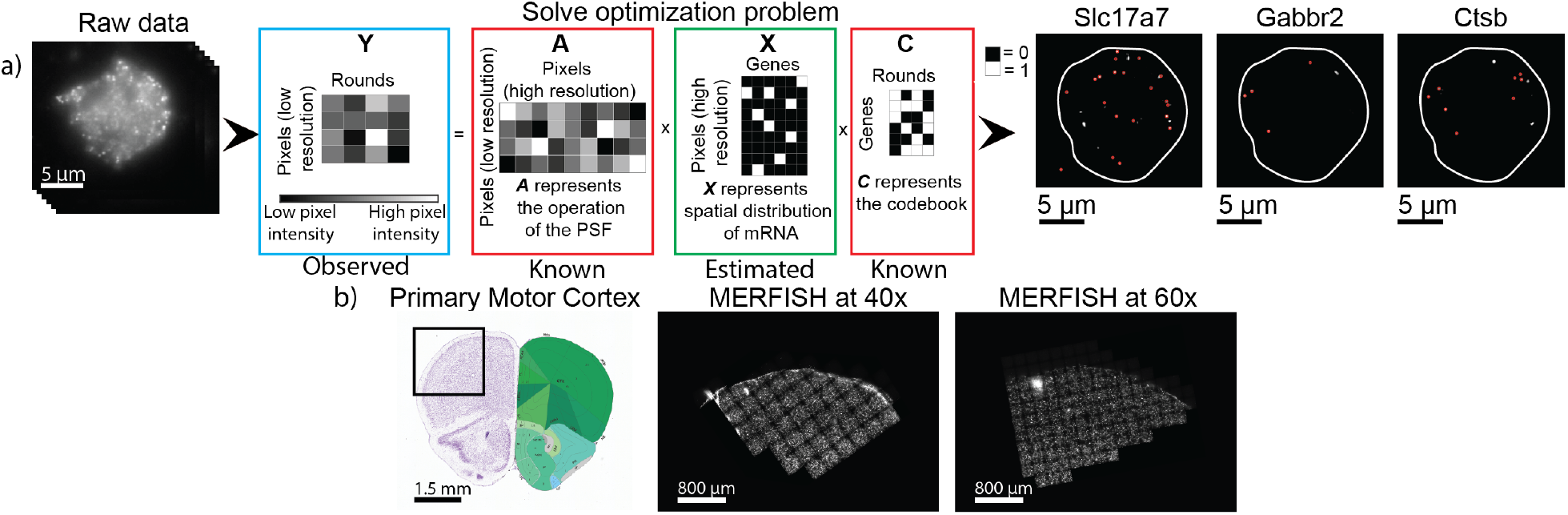
Summary of JSIT. a) Schematic of JSIT pipeline. Left: MERFISH signal is acquired through multiple rounds of imaging. Center: MERFISH data is modeled as a matrix product. With this framing, decoding is a sparse recovery problem. Right: JSIT produces maps of the distribution of mRNA transcripts of each gene. b) Schematic of experiment. Left: Allen Brain Atlas depiction of the mouse brain, with cell regions labeled on right. Primary motor cortex (MOp) highlighted. (Lein et al. 2007) Center: Downsampled MERFISH image of MOp, imaged at 40x. Right: Downsampled MERFISH image of MOp, imaged at 60x.

To better resolve spots in the acquired data, we define the matrix 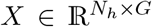 at a higher resolution than *Y*, i.e. *N*_*h*_ *> N*_*l*_. Specifically, we pick a scale factor *s*, and divide each pixel represented in *Y* into *s*^2^ equally-sized sub-pixels, giving *N*_*h*_ = *s*^2^*n*^2^ = *s*^2^*N*_*l*_. The nonzero values of elements of *X* indicate the intensity of fluorophores bound to mRNA molecules, while zeros indicate no mRNA present. Next, we define the PSF matrix 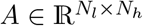, such that element (*i*_*l*_, *i*_*h*_) represents the percentage of photons originating at location *i*_*h*_ in the sample distributed to pixel *i*_*l*_ in the image; *A* can be obtained theoretically or empirically for any optical system used to image the sample. Finally, we define the codebook (*g, f*) of *C* by setting element 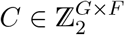 equal to the value of bit *f* of barcode *g*. For any iST experimental protocol, *C* is known. Our goal is to recover *X* from *Y*.

### 2.2 Joint Sparse method for Imaging Transcriptomics (JSIT)

#### 2.2.1 Preliminary denoising

To minimize the contribution of the noise term *η*, we filter both high- and low-frequency noise by convolving each of the *F* observed images with a difference-of-Gaussians band-pass filter, where one Gaussian is empirically chosen to be broader than the microscope PSF and the other narrower. In processing the 60x MERFISH data, for example, we used the filter:

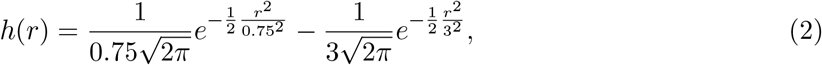

where 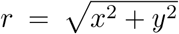, with *x* and *y* representing distance in pixels. The resulting images are vectorized and concatenated to form 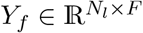.

### 2.3 Decoding as a sparse recovery problem

We seek to recover *X* from *Y*_*f*_ by solving a regularized least-squares optimization problem, using principles of compressed sensing (Eldar and Kutyniok 2012; Eldar 2015). We impose several constraints: a) only one molecule is present at any given location, so the rows of *X* are one-sparse; b) molecules are present at few locations in the FOV, so *X* is row-sparse; c) the FOV contains few mRNA molecules, so *X* is overall sparse. We choose to impose constraint (a) in post-processing rather than as part of the optimization problem, finding that this improves results, similar to (Mazor et al. 2018). We impose (b) with a mixed *ℓ*_1_*/ℓ*_2_ norm on the rows of *X*, and (c) by an *ℓ*_1_ norm on *X*. These functions are combined convexly, as in the Sparse Group LASSO formulation (Simon et al. 2013), leading to the optimization problem:

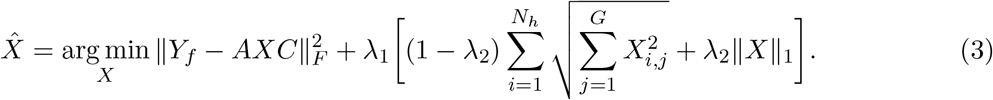

We solve (3) with the Fast Iterative Shrinkage-Thresholding Algorithm (FISTA) (Beck and Teboulle 2009; Eldar and Kutyniok 2012), using the proximal operator for the regularization function as given in (Bach et al. 2012). These are both detailed in supplementary Section S3.1.

After obtaining 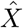, one-sparsity is imposed on the rows of 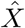 by setting to 0 all but the maximal value of each row of *X*, rejecting small elements representing spuriously detected molecules. The resulting 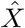 is additionally hard-thresholded as a de-noising mechanism. Having recovered 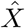, each column *i* of 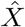 is reshaped as an image, giving the spatial distribution of gene *i*. Clusters of nonzero elements are formed by identifying connected components using the Matlab command `bwconncomp` and the centroid of each cluster is taken to be the location of a molecule associated with gene *i*. This addresses cases where each molecule is represented by multiple nonzero elements in 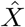, due to small variations in the spot size, as can be seen in Fig. 1a. We reject clusters of pixels below an empirically chosen area threshold, to minimize spurious calls.

The resulting method, called the Joint Sparse method for Imaging Transcriptomics (JSIT), is summarized in Algorithm 1. For comparison, the MERlin decoding procedure is detailed in supplementary Section S2.

#### Algorithm 1

JSIT

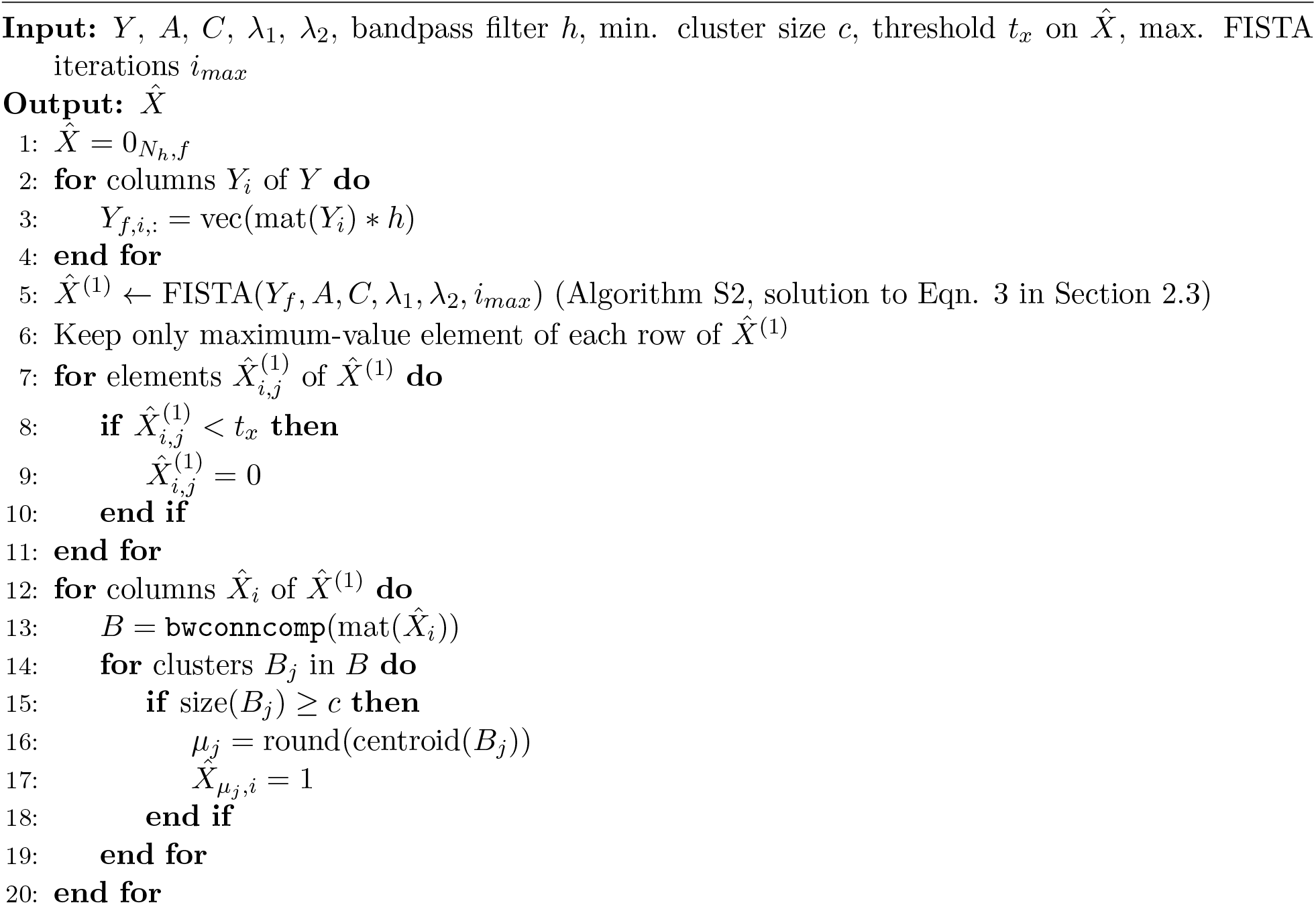

#### 2.3.1 Post processing and data cleanup

After the main decoding step, false positive detections are minimized by adaptively filtering decoded spots. We use a method very similar to MERlin (Moffitt, Hao, et al. 2016), described in Supplementary Section S3. Decoded transcripts are then assigned to individual cells using a seeded watershed approach (Meyer 1994). We reject cells with area below one-half of the median area or above two times the median area. Transcripts that fall within the boundaries generated by the segmentation are then assigned to cells, to construct a count table 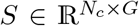, where *N*_*c*_ is the number of cells in the ROI, and element *S*_*i,j*_ is the number of decoded transcripts of gene *j* within the bounds of cell *i*. The centroids of the segmented areas of each cell are also obtained, and a table of locations is created, 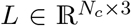, with the *i*th row of *L* a vector giving the 3-dimensional position in the ROI of the *i*th cell. We also produce a table of “relative” locations, 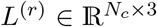, in which the *i*th row is a vector giving the position within its FOV of the *i*th cell.

Finally, we correct for observed expression differences between cells at the center and edges of the FOV, presumably caused by spherical aberrations of the microscope objective affecting imaging and decoding performance. We bin cells in all FOVs based on distance to the FOV center into *K* bins, and calculate the average expression level *m*_*g,k*_ for each of *G* genes, and scale the abundance of each gene in each cell by multiplying each element of *S* by 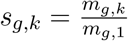, according to the bin *k* to which its cell belongs.

#### 2.3.2 Parameter tuning

Three parameters require tuning in the JSIT workflow: the regularization parameter *λ*_1_, the threshold for *X, t*_*x*_, and the minimum accepted cluster size *t*_*c*_ (*λ*_2_ can also be tuned, but here is set to 0.5 throughout). The parameter *λ*_1_ controls the sparsity level of the results: as *λ*_1_ is increased, more values of *X* are equal to zero, and, fewer spots are be decoded. However, the adaptive filtering step complicates this: as *λ*_1_ is decreased, the number of spots decoded increases, and, along with this, the number of spots corresponding to blank barcodes increases. This can result in an increased number of coding barcodes being rejected, and a decrease in the total number of postadaptive-filtering decoded spots, even as the total number of pre-adaptive-filtering decoded spots increases. We found that the best spatial homogeneity results (see Section 2.4.2) are obtained when the number of post-adaptive-filtering decoded spots is maximal, and so we use this as a heuristic for selection of *λ*_1_. To select *λ*_1_, we decode a small number of FOVs with various values of *λ*_1_, perform adaptive filtering, and select the value which gives the highest number of post-filtering spots (Supp. Fig. S2). After selecting *λ*_1_, we use the same method to select *t*_*x*_. We also find that performance of the pipeline improves when clusters below a certain size are rejected before adaptive filtering. Precision (% of detected molecules corresponding to real molecules) increases with the minimum cluster size *t*_*c*_, as many small clusters represent noise, while recall (% of molecules detected) decreases as the minimal cluster size increases, because some real molecules are represented by small spots. We typically set *t*_*c*_ to 2.

### 2.4 Validation of JSIT

#### 2.4.1 Dataset

We obtained two sequential coronal slices of mouse MOp (separated by 10 μm), and prepared them for MERFISH imaging according to the procedure described in (Moffitt, Hao, et al. 2016) (Fig. 1b). We probed library of 115 genes, designed for cell typing in the cortex the mouse brain (Stogsdill et al. 2022). While this library was specifically designed to study the primary somatosensory cortex (SSp), the SSp has similar cell type structure to the MOp, and the library includes marker genes for the major cell types present in the MOp. Readout and encoding probes were obtained from Vizgen Inc., and sequential slides were imaged using Vizgen’s MERSCOPE alpha instrument. One slice was imaged using a 60x NA 1.4 objective, and the other was imaged using a 40x NA 0.95 objective. We captured 168 x-y locations covering 6.72 mm^2^ and roughly 6000 cells at 60x and 51 x-y locations covering 4.59 mm^2^ and roughly 4500 cells at 40x. In both datasets we acquired seven z-positions per x-y location, separated by 1.5 μm. Illumination intensities and exposure times were kept the same in each dataset.

#### 2.4.2 Cell cluster and localization analysis

After producing the cell-by-gene count table *S*, statistical results were computed on an aggregate level, calculating the average number of transcripts decoded per cell, the average number of genes with nonzero numbers of transcripts per cell, and the average intensity of a detected transcript (throughout the text, these results are expressed as a mean ± standard deviation unless otherwise noted. Statistical tests and *p* values are described next to individual results, with results treated as significant if *p <* 0.05, unless otherwise stated). Then, standard single-cell analysis techniques were used to cluster cells by their gene expression levels. All analysis was performed using the Scanpy library in Python (Wolf, Angerer, and Theis 2018). The count table was first normalized such that each cell would have the same number of total counts, and dimensionality reduction was performed using principle component analysis (PCA), keeping the top 50 principle components. We clustered the cells in principle component space using the Louvain clustering algorithm (Blondel et al. 2008), selecting a resolution parameter by empirically adjusting until the major cortical cell types were revealed: excitatory neurons, interneurons, microglia, oligodendrocytes, and astrocytes. Following this analysis, we removed *Slc17a7*, which was highly expressed in all excitatory neurons and repeated these steps to subcluster excitatory neurons to reveal cortical layer specific subpopulations. We identified spatial regions belonging to major layers of the primary motor cortex—Layers 2/3, 4, 5, and 6—by visualizing the spatial expression of well-known marker genes of each layer (*Stard8, Rorb, Deptor*, and *Tle4*), and manually annotating the areas of high expression, as shown in Fig. 3c. To compare the cluster assignments in gene-expression space to the spatial layer assignments, we used the cluster homogeneity score (Rosenberg and Hirschberg 2007), defined as:

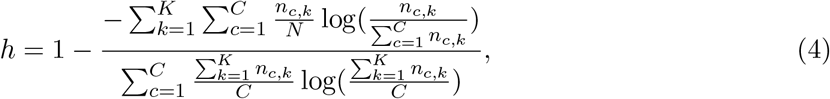

where *K* is the number of spatial layers, *C* is the number of gene-expression space clusters, and *n*_*c,k*_ is the number of cells belonging to cluster *c* and layer *k*. This metric quantifies the notion that excitatory neurons assigned to the same cluster in gene-expression space should belong to the same spatial layer. This metric can obtain a high score even if a layer is comprised of multiple gene-expression space clusters, each confined to that layer. This is biologically reasonable as layers may be subdivided into multiple cell types. The homogeneity score is related to the ratio of the conditional entropy of layers given cluster assignments to the overall entropy of the layers. The score is between 0 and 1, with 1 signifying perfect homogeneity.

## 3 Results

We benchmarked the performance of the JSIT and MERlin pipelines by analyzing MERFISH data from sequential slices of the mouse MOp, acquired with both 60x and 40x magnification. First, we focused on processing the conventional, high-magnification (60x) data with JSIT and MERlin, comparing the two pipelines at the level of individual mRNA transcript decoding, measuring aggregated results like the average number of molecules per cell, and qualitatively comparing the spatial distributions of the expression of individual genes to the distributions seen in the Allen Brain Atlas (ABA)(Lein et al. 2007). Then, at the level of the spatial organization of cell types, we used a metric called spatial homogeneity, defined above in section 2.4.2, to quantify the degree to which cell types known to be spatially confined (identified by unsupervised clustering on gene expression) align with spatial layers. We then performed the same suite of analyses on the data acquired at 40x, and investigated the differences between the approaches in overall throughput, considering both imaging and computation time.

On aggregate measures of transcripts, there was good agreement between JSIT and MERlin results. We first measured pseudo-bulk abundances of the 115 profiled genes in 60x data, and found high gene expression correlation between JSIT and MERlin (Pearson’s r=83%). MERlin detected a significantly larger number of transcripts per cell relative to JSIT (587± 342, *n* = 6068 cells vs 400± 291, *n* = 5809 cells, *p* = 6.7 × 10^*−*217^, Welch’s t-test), but MERlin and JSIT did not detect a significantly different number of genes with nonzero counts per cell (61± 18, n=6068 vs 60± 22, *n* = 5809, *p* = 0.329, Welch’s t-test) (Fig. 2a). Because ground truth data is difficult to acquire in parallel with MERFISH, it is not clear whether the additional calls by MERlin represent true or false positives. To understand the apparent higher sensitivity of MERlin in comparison to JSIT, we thus calculated the average pixel intensity of the signal from decoded transcripts, finding that the average pixel intensity for MERlin calls was lower than that of JSIT calls (19±15 post-denoising counts, *n* = 8.4 × 10^6^ vs. 40±16, *n* = 4.5 × 10^6^, *p <* 2.2 × 10^*−*308^, Welch’s t-test) (Fig. 2a). This was also apparent qualitatively: when examining the raw data for individual transcript calls, JSIT’s calls generally are brighter than MERlin’s (Supp. Fig. S1) The additional, dimmer spots identified by MERlin may include more spurious calls, as autofluorescence noise frequently appears as dim spots.

**Figure 2:**
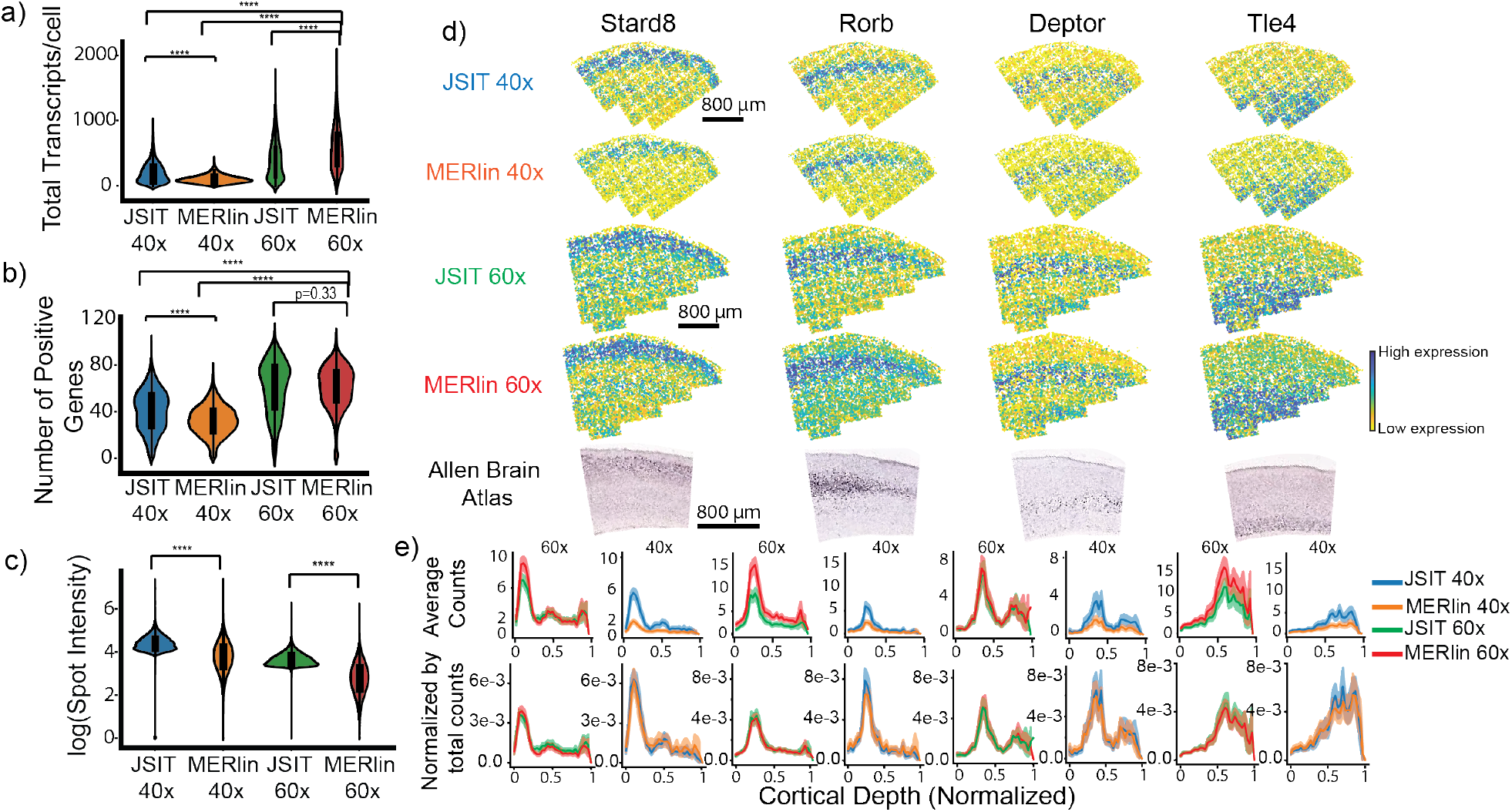
JSIT accurately recovers gene expressions. a)-c) Transcript quantification in decoded results, by decoding pipeline. Asterisks indicate that differences between distributions are statistically significant. a) Number of transcripts per cell. b) Total genes with nonzero counts per cell. c) Intensity of decoded molecules. d) Spatial distribution of several genes with known spatial patterns as decoded by MERlin and JSIT from 40x and 60x data, and spatial distribution of genes as given by the Allen Brain Atlas (ABA) (Lein et al. 2007). e) Spatial distribution of expression of spatially varying genes versus cortical depth. Top row: Average number of mRNA molecules per cell, in each depth bin. Bottom row: Average number of mRNA molecules per cell, normalized by total counts.

**Figure 3:**
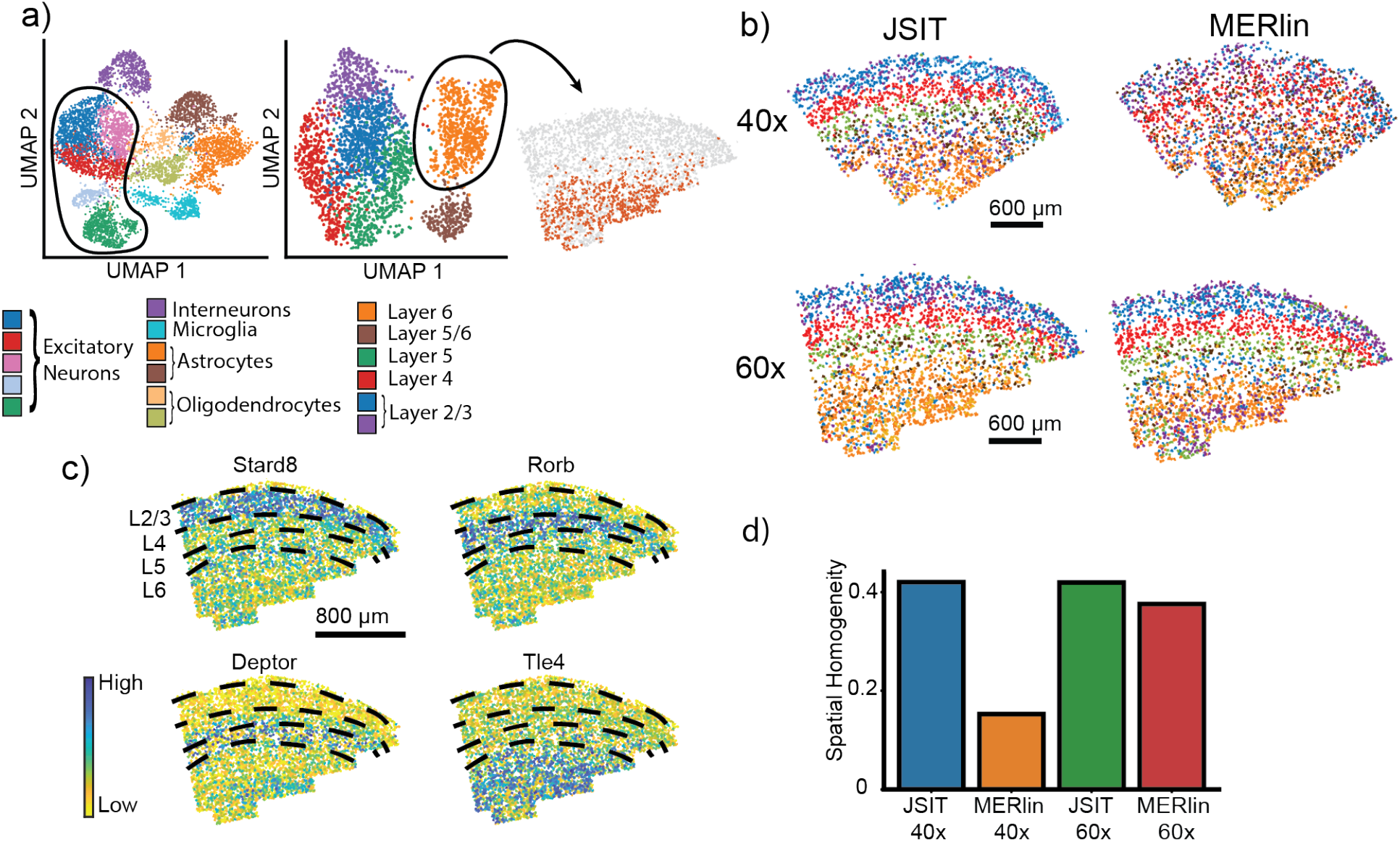
JSIT accurately recovers cortical layers of excitatory neurons. a) UMAP plots of the gene expression cells decoded by JSIT, imaged at 60x magnification. Left: all cells, with clusters associated with excitatory neurons outlined. Center: re-clustering of excitatory neurons. Right: Mapping of one cluster (L6 excitatory neurons) to spatial coordinates. b) Spatial distribution of cell-types identified by unsupervised clustering of excitatory neurons in results from JSIT and MERlin, on 40x and 60x data. c) Defined spatial layers of excitatory neurons in the cortex (2/3, 4, 5, and 6), in 60x data, overlaid on distribution of key marker genes as detected in JSIT 60x data. d) Spatial homogeneity is computed between cell type clusters and spatial layers shown in c).

We then examined the spatial distribution of the expression of several marker genes of cortical layers in the MOp (*Stard8, Rorb, Deptor*, and *Tle4*), finding that in both MERlin and JSIT, they qualitatively matched those recorded in the ABA (Lein et al. 2007) (Fig. 2b). When we measured the distribution of these genes as a function of cortical depth, and normalized by the total number of transcripts detected, JSIT and MERlin produced nearly identical distributions, though MERlin showed higher transcript numbers overall (Fig 2c). We conclude that on 60x data, JSIT provides similar bulk level performance to MERlin.

Next, we performed higher-level analysis, comparing the results of clustering data processed by JSIT and MERlin by gene expression. It is well-known from scRNA-seq data that cortical excitatory neurons in different layers of the mouse brain have distinct expression profiles (Zhang et al. 2021). Thus, unsupervised clustering of neurons by gene expression should result in clusters corresponding to layer-specific subtypes. When we performed this analysis, we found that for both MERlin and JSIT, the cell type clusters mapped to regions lying within the well-known cortical layer boundaries (Fig. 3a). We also computed a metric called spatial homogeneity, described in Section 2.4.2, which quantifies the extent to which observations that fall into the same class in the clustering by gene expression also fall into the same spatial layer. We found JSIT produced clusters with spatial homogeneity of 0.41, higher than the spatial homogeneity of 0.38 achieved by MERlin (Fig. 3d), showing that JSIT reproduces the laminar spatial structure of cortical neurons at least as well as MERlin. As a whole, these results showed that, with 60x high-magnification data, the results produced by JSIT were very similar to those produced by MERlin, with slightly lower sensitivity.

We next performed the same set of analyses on the results produced by processing the 40x data with MERlin and JSIT, starting with transcript-level aggregate metrics. Pseudo-bulk correlation with the MERlin 60x data was high for both JSIT 40x (Pearson’s r=0.94) and MERlin 40x (Pearson’s r=0.97). MERlin and JSIT both detected significantly fewer transcripts per cell at 40x than MERlin detects at 60x (MERlin 40x: 104± 60, *n* = 4323, JSIT 40x: 200± 144, *n* = 4288, Welch’s t-test gives *p <* 2.2 × 10^*−*308^ for both MERlin 40x and JSIT 40x in comparison to MERlin 60x), but JSIT 40x detects significantly more transcripts per cell than MERlin 40x (*p <* 2.3 × 10^*−*308^, Welch’s t-test). The same pattern is seen in computing mean genes with nonzero counts per cell, with both MERlin 40x and JSIT 40x detecting significantly fewer positive genes per cell than MERlin 60x (MERlin 40x: 32± 13, JSIT 40x: 41± 17, *p <* 2.2 × 10^*−*308^, Welch’s t-test for both in comparison to MERlin 60x), with JSIT 40x detecting significantly more positive genes per cell than MERlin 40x (*p* = 1.1 × 10^*−*139^, Welch’s t-test). In computing the average intensity of decoded spots, we again found that JSIT calls were brighter than MERlin spots (87±43 post-filtering counts, *n* = 1.6 × 10^6^ vs. 53±41 *n* = 1.2 × 10^6^, *p <* 2.2 × 10^*−*308^, Welch’s t-test) (Fig. 2a). We conclude that JSIT’s 40x results were closer than MERlin’s 40x results to the results of JSIT or MERlin at 60x, although both MERlin and JSIT produced less-sensitive results at 40x magnification.

When comparing spatial patterns of gene expression, the JSIT 40x results look qualitatively similar to the 60x JSIT results and the ABA, while the results of using MERlin show sparser expression patterns, especially for *Stard8* (Fig. 2b). As a function of cortical depth, we observed the same pattern: for each gene, JSIT detected substantially more transcripts in the region for which the gene is a marker, and the normalized frequency distribution generated by the JSIT had a higher peak and lower trough than that generated by MERlin (Fig. 2c).

While noting that 40x data analyzed by JSIT had lower detection sensitivity than 60x data, the qualitative similarity between the MERlin 60x and JSIT 40x gene distributions led us to consider whether we could still use JSIT 40x data to perform the types of cell-based downstream analyses that iST typically targets. We thus clustered the cells of both the JSIT 40x and MERlin 40x count tables by gene expression and computed the spatial homogeneity of sub-clustered excitatory neurons. The gene expression clusters from JSIT map well to the cortical layers, as defined by expression of marker genes (Fig. 3c). The gene expression clusters from MERlin are much less spatially confined (Fig. 3b). The JSIT 40x results have spatial homogeneity higher than even the MERlin 60x results, 0.40, while the MERlin 40x results have much lower spatial homogeneity, 0.18 (Fig. 3d). So, while JSIT may exhibit lower sensitivity when decoding 40x data, it is nonetheless able to produce expression clusters that match the laminar structure in this data as well as MERlin on 60x data.

To test the similarity of the cell-type clusters found by JSIT 40x and MERlin 60x we clustered the JSIT 40x and MERlin 60x results together. Despite some batch effects, we noted that all but two clusters (which correspond to the JSIT 40x and MERlin 60x clusters for L2/3 excitatory neurons) contained many cells from JSIT 40x and MERlin 60x (Fig. 4a), showing that we obtain the same major cell types in the results from both JSIT 40x and MERlin 60x. Additionally, we found that clusters labeled as L4, L5, and L6 excitatory neurons mapped to the same spatially confined layers in the JSIT 40x and MERlin 60x data (Fig. 4b). This confirmed to us JSIT’s ability to reproduce the clustering results of MERlin 60x at 40x magnification, enabling higher throughput.

**Figure 4:**
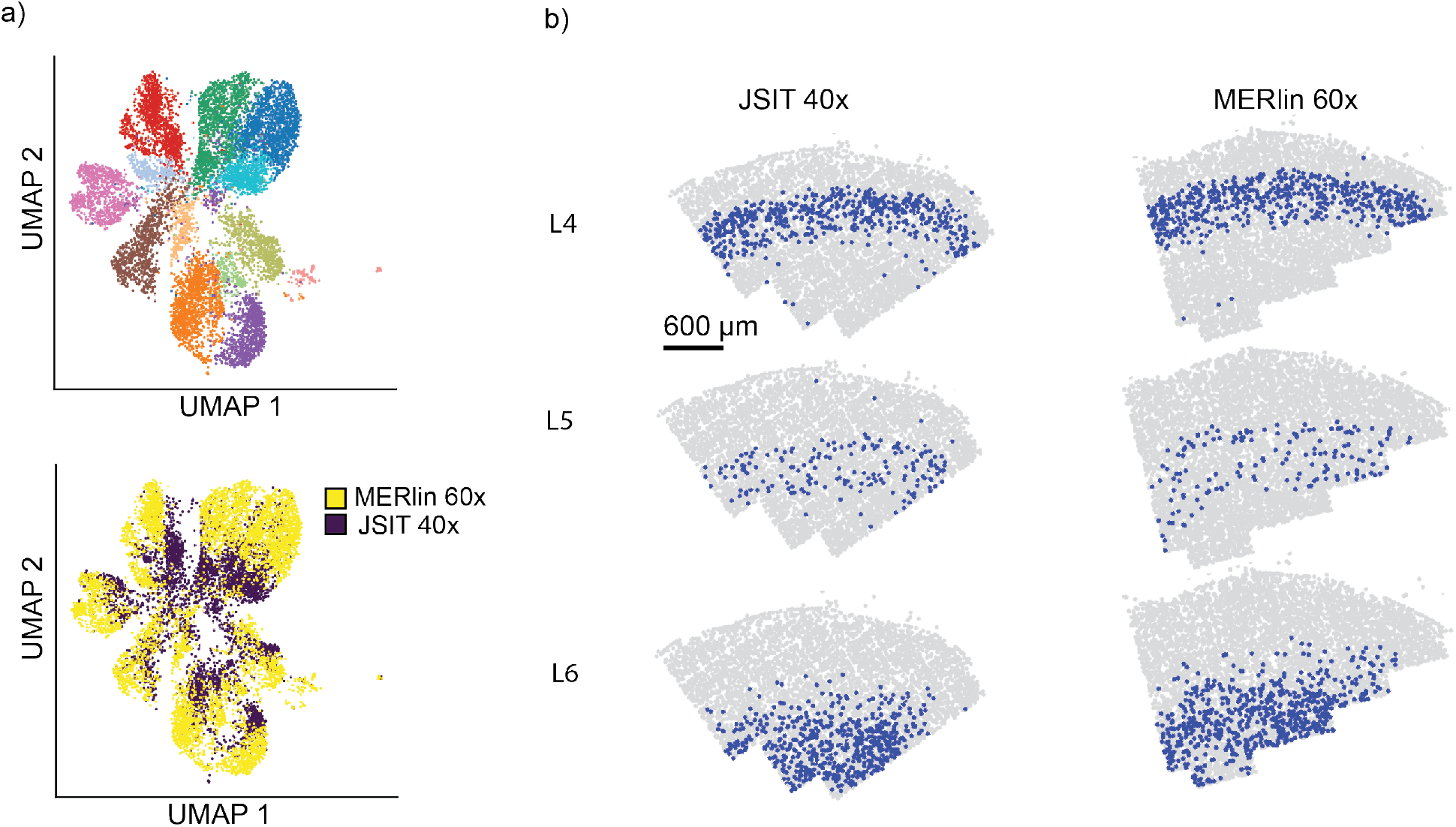
JSIT 40x clustered with MERlin 60x. a) UMAP, and clustering with Louvain clustering algorithm, of JSIT 40x results together with MERlin 60x results. b) Spatial mapping of cells clustered as excitatory neurons, when JSIT 40x and MERlin 60x results are clustered together.

In all, we found that JSIT enables accurate cell typing of MERFISH when imaging at 40x, with somewhat lower sensitivity than when imaging at 60x. In the common scenario in which the priority is accurate cell typing, and where parallelized computational resources are relatively available, JSIT enables increased throughput. Thus, JSIT allows the profiling of larger volumes of tissue and allow transcriptomal study of broader spatial contexts, which is necessary to better understand larger-scale systems of genetic behavior.

## 4 Discussion

We have presented a new algorithmic pipeline, JSIT, for decoding multiplexed iST data. JSIT uses an optimization-based method to take advantage of the sparse nature of IT signal, and, with high-magnification imaging, shows comparable ability to detect gene expression, and perform cell typing, in comparison to the currently used decoding pipeline, MERlin.

We found that JSIT decoded MERFISH data acquired using a 40x microscope objective with higher sensitivity than MERlin on the same data, although lower sensitivity than MERlin’s decoding of 60x data. Despite this drop of sensitivity relative to 60x data, when the gene count matrix obtained by decoding 40x data with JSIT is clustered by gene expression using standard pipelines, clusters corresponding to all major cell types are correctly identified, with higher spatial homogeneity than that obtained by using MERlin to decode 60x data. In contrast, the gene count matrix obtained by decoding 40x data with MERlin was unable to produce cell-type clusters which correctly correspond to cell types, especially in excitatory neurons. We conclude that decoding with JSIT enables accurate cell typing while imaging at lower magnification.

Given the scale of iST data (frequently terabytes), it is important to consider the computational resources required to implement JSIT. Because the JSIT algorithm involves many large matrix multiplications, it requires substantially more computational time per FOV than MERlin: almost 67 fold more for the same area. While this is a significant difference, we note that MERlin performance has been optimized several times, while this is the first presentation of JSIT; that computational load is easier to parallelize than experimental load; and that computational costs for MERFISH experiments are dominated by storage rather than by compute. In our cost models, the decreased storage costs of lowering the number of FOVs stored by 2.25x pays for the difference in compute time expense in the first year of the project. So, in cases in which computational resources are readily available, and in which the key priority is identification of cell types, JSIT can be used to study larger tissue samples in less time, enabling a greater range of insights into relationships between different cell types.

While in this study we have applied JSIT only to MERFISH data, the optimization-based method may be applied to any multiplexed IT method, such as seqFISH+ (C. Eng et al. 2019), or STARmap (Wang et al. 2018). We suggest that JSIT or an adapted version be applied widely to other IT strategies. Finally, JSIT is readily extensible due to its straightforward model of iST signal generation: more complex noise models may be incorporated, for example, or JSIT may be extended to a 3D analysis of IT data sampled more densely by depth. The optimization-based approach also opens the door to the incorporation of machine learning via algorithm unrolling (Monga, Y. Li, and Eldar 2021; Sahel et al. 2022), as in (Dardikman-Yoffe and Eldar 2020), which can also reduce computational costs. Extensions are likely to improve the performance of JSIT and improve ability to study the spatial distributions of gene expression.

## Acknowledgements

We thank Mehrtash Babadi, Luca d’Alessio, Gili Dardikman-Yoffe, and Yair Ben Sahel for helpful discussions.

## Data Availability

Software implementation of JSIT, together with example files, are available at https://github.com/jpbryan13/JSIT. Raw MERFISH data is available from the authors on request.

