## Supplemental Information for "Optimization-Based Decoding of Imaging Spatial Transcriptomics Data"

### Supplementary Material

**Contents**

**S1** Detailed MERlin decoding algorithm

**S2** Adaptive filtering to remove false positives

**S3** FISTA algorithm to solve Eqn. 3 (Section 2.3)

**S4** Supplemental Figures

- **Figure S1** Individual spot decoding from raw MERFISH data
- **Figure S2** Parameter selection of $\lambda$ and $t_x$

### S1 MERlin

**Algorithm S1** MERlin decoding

**Input:** $Y$, $C$, bandpass filter $h_b$, PSF estimate $h_p$, decoding threshold $t_d$, min. cluster size $c$
**Output:** $\hat{X}$

$\hat{X} = 0_{N_h,f}$
**for** columns $Y_i$ of $Y$ **do**
  $Y_{f,i,:} = \text{vec}(\text{mat}(Y_i) * h_b)$
  $Y_{f,i,:} = \text{RL}(Y_{f,i,:}, h_p)$ (Richardson-Lucy deconvolution (Richardson 1972))
**end for**
**for** rows $Y_j$ of $Y$ **do**
  $d = \min_i \|C_{i,:} - Y_j\|^2$
  $\hat{i} = \arg\min_i \|C_{i,:} - Y_j\|^2$
  **if** $d < t_d$ **then**
    $\hat{X}_{j,i} = 1$
  **end if**
**end for**
**for** columns $\hat{X}_i$ of $\hat{X}^{(1)}$ **do**
  $B = \texttt{bwconncomp}(\text{mat}(\hat{X}_i))$
  **for** clusters $B_j$ in $B$ **do**
    **if** $\text{size}(B_j) \geq c$ **then**
      $\mu_j = \text{round}(\text{centroid}(B_j))$
      $\hat{X}_{\mu_j,i} = 1$
    **end if**
  **end for**
**end for**

### S2 Adaptive Filtering

To control for false positives, within each codebook several barcodes exist which are not assigned to any of the targeted genes. We bin the spots by the average value of the elements of $\boldsymbol{X}$ corresponding to the cluster, the cluster size and the average norm of the pixel traces, and for each bin compute the percentage of called spots which are assigned to blank barcodes. Then, the secant method is used to find a threshold on the acceptable percentage of blank barcodes which, when applied to the histogram, a specific misidentification rate of 0.05. Spots belonging to bins with a fraction of blank barcodes below this threshold are dropped from further analysis. Given a codebook with $N_b$ blank barcodes and $N_c$ coding (i.e. not blank) barcodes, and a dataset with $B$ spots assigned to blank barcodes and $C$ spots assigned to coding barcodes, misidentification rate is defined as:

$$m = \frac{\frac{B}{N_b}}{\frac{C}{N_c}}. \tag{1}$$

.

### S3 FISTA for JSIT

**Algorithm S2** FISTA solution of Eqn. 3 (Section 2.3)

**Input:** $Y$, $A$, $C$, $\lambda_1$, $\lambda_2$, $i_{max}$
**Output:** $\hat{X}^{(k)}$

$K = CC^T$
$M = AA^T$
$[U_k, S_k, V_k] = \text{SVD}(K)$
$[U_m, S_m, V_m] = \text{SVD}(M)$
$L_f = S_{k,1,1} S_{m,1,1}$
$\hat{X}^{(k)} = 0$
$Z = 0$
$t = 1$
$i = 1$
**while** $i \leq i_{max}$ **do**
    $G = A^T(AZC - Y)C^T$
    $X_p \leftarrow \hat{X}^{(k)}$
    $\hat{X}^{(k)} = \text{prox}_{\text{SGL}}(\frac{\lambda}{L_f}, \lambda_2, \hat{X}^{(k)} - \frac{G}{L_f})$ (Algorithm S3)
    $t_p \leftarrow t$
    $t \leftarrow \frac{1+\sqrt{1+4t^2}}{2}$
    $Z \leftarrow X^{(k)} + \frac{t}{t_p}(X^{(k)} - X_p)$
    $i \leftarrow i + 1$
**end while**

**Algorithm S3** Proximal of Sparse Group Lasso

**Input:** $\alpha$, $\beta$, $X$
**Output:** $\text{prox}_{\text{SGL}}(\alpha, \beta, X)$

$P = \max(|X| - \alpha\beta)\text{sgn}(X)$
**for** rows $P_i$ of $P$ **do**
    **if** $\|P_i\| \geq (1-\alpha)\beta)$ **then**
        $P_i = \frac{1-(1-\alpha)\beta}{\|P_i\|} P_i$
    **end if**
**end for**
$\text{prox}_{\text{SGL}}(\alpha, \beta, X) = P$

#### S4 Supplemental Figures

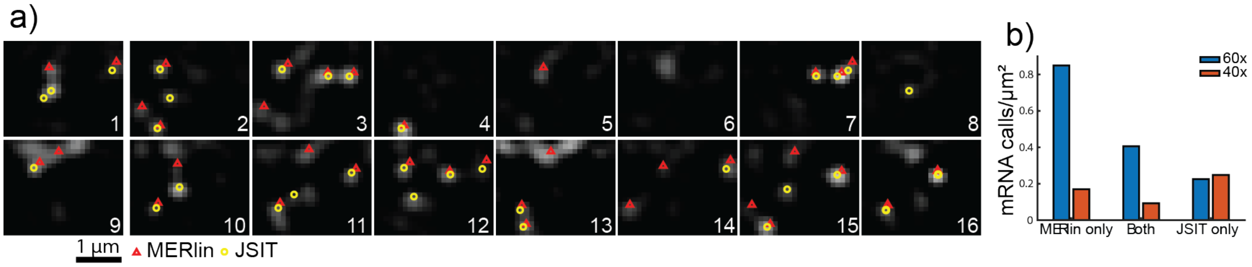


Figure S1: a) Imaging data from each of 16 frames of 60x MERFISH data, overlaid with estimates by MERlin and JSIT of positions of transcripts. b) Bar plot showing the number of transcripts per µm$^2$ identified by only MERlin, only JSIT, and by both pipelines, at both 60x and 40x.

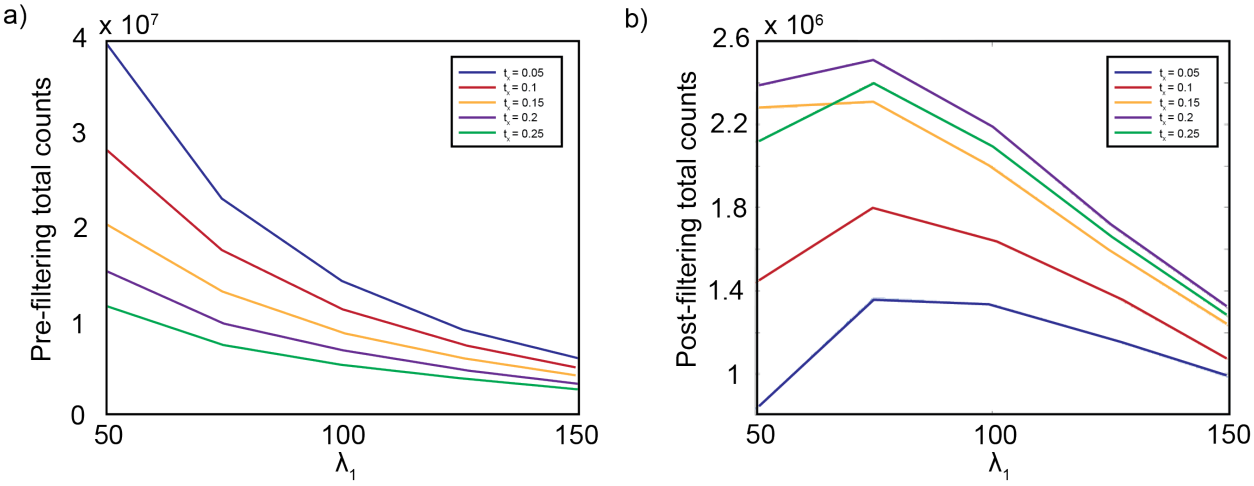


Figure S2: **Parameter selection of $\lambda_1$ and $t_x$** a) As $\lambda_1$ increases, total counts before adaptive filtering decreases. As $t_x$ increases, total counts decrease. b) Below a certain point, decreasing $\lambda_1$ causes total counts after adaptive filtering to decrease. The same is true of $t_x$.
